# Dynamic metabolic network modeling of a mammalian cell cycle using time-course multi-omics data

**DOI:** 10.1101/2021.10.11.464012

**Authors:** Ho-Joon Lee, Fangzhou Shen, Alec Eames, Mark P. Jedrychowski, Sriram Chandrasekaran

**Affiliations:** Department of Genetics; Yale Center for Genome Analysis, Yale University, New Haven, USA; Department of Biomedical Engineering, University of Michigan, Ann Arbor, USA; Department of Cell Biology, Harvard Medical School, Boston, USA; Department of Biomedical Engineering; Michigan Institute for Data Science; Rogel Cancer Center; Center for Computational Medicine and Bioinformatics, University of Michigan, Ann Arbor, USA

## Abstract

Cell cycle is a fundamental process for cell growth and proliferation, and its dysregulation leads to many diseases. How metabolic networks are regulated and rewired during the cell cycle is unknown. Here we apply a dynamic genome-scale metabolic modeling framework (DFA) to simulate a cell cycle of cytokine-activated murine pro-B cells. Phase-specific reaction activity predicted by DFA using time-course metabolomics were validated using matched time-course proteomics and phospho-proteomics data. Our model correctly predicted changes in methionine metabolism at the G1/S transition and the activation of lysine metabolism, nucleotides synthesis, fatty acid elongation and heme biosynthesis at the critical G0/G1 transition into cell growth and proliferation. Metabolic fluxes predicted from proteomics and phosphoproteomics constrained metabolic models were highly consistent with DFA fluxes and revealed that most reaction fluxes are regulated indirectly. Our model can help predict the impact of changes in nutrients, enzymes, or regulators on this critical cellular process.

## INTRODUCTION

The cell cycle is a fundamental biological process vital for growth and survival of organisms. Many proteins and genes that are involved in the cell cycle regulation and control have been identified and characterized across diverse species (Hartwell and Weinert, 1989; Hunt, 1991; Hunt et al., 2011; Norbury and Nurse, 1992; Nurse, 1994; Nurse, 2000). Each cell cycle in healthy cells is typically characterized by 4 distinct phases under precise control: G1 phase for cell growth and molecular preparation for DNA synthesis, S phase for DNA synthesis, G2 phase for molecular preparation for cell division, and M phase for cell division. If cells are not dividing or proliferating, they are in a resting or G0 state. Dysregulation of the cell cycle often leads to diseases such as cancer, diabetes, heart disease, neurodegenerative disease, and aging (Ahuja et al., 2007; Bicknell and Brooks, 2008; Boehm and Nabel, 2003; Chandler and Peters, 2013; Cozar-Castellano et al., 2006; El-Badawy and El-Badri, 2016; Hartwell, 1992; Hartwell et al., 2006; Hartwell et al., 1994; Hartwell and Kastan, 1994; Hinault et al., 2008; Icard et al., 2019; Kamb, 1995; Massague, 2004; Rane and Reddy, 2000; Sarsour et al., 2009; Swanton, 2004).

The cell cycle regulation in mammalian cells is more complex than that in simpler organisms such as yeast. With advent of high-throughput sequencing technologies, several transcriptomic analyses have been performed in mammalian cells to study cell cycle genes on a global level (Bar-Joseph et al., 2008; Grant et al., 2013; Liu et al., 2017; Pena-Diaz et al., 2013; Whitfield et al., 2002). The numbers of cell cycle genes identified in those studies vary from ~650 to ~2500. In contrast to transcriptomics, studies on proteomics and especially metabolomics of cell cycle have been generally lacking. To address this gap, we previously carried out mass spectrometrybased quantitative temporal proteomics and metabolomics to study an IL-3-dependent cell cycle in murine pro-B cells (Lee et al., 2017). We found that the cell cycle in murine pro-B cells exhibit cancer-like metabolic reprogramming at multiple molecular levels. In addition, the methionine consumption rate is the highest in the G1 phase among all essential amino acids.

Our data also highlight the complexity of metabolic changes that occur during the cell cycle. A theoretical model that explains the dynamics of this process is currently lacking. Such a model would allow us to understand the ramifications of alterations in the levels of enzymes, nutrients, metabolites, or regulators in this critical cellular process. However, it is challenging to create such a model as a typical mammalian cell contains thousands of metabolic reactions. This metabolic network is in turn regulated by an equally complex network of regulators, including transcription factors for gene expression and protein kinases for phosphorylation. The size and complexity of metabolism and its regulation makes it challenging to predict the metabolic behavior of cells.

To address this, here we apply a theoretical framework called Genome-Scale Metabolic-modeling (GSM). GSM is widely used to study metabolism of systems with thousands of reactions and to predict the metabolic state of various organisms based on thermodynamics, mass-balance and stoichiometric constraints (Nilsson and Nielsen, 2017; O’Brien et al., 2015; Shlomi et al., 2008; Yizhak et al., 2015). GSM can calculate metabolic reaction activity (i.e., fluxes) on a global level, which is infeasible to measure experimentally.

GSM methods assume a steady state of metabolite concentration and hence not directly applicable to dynamic systems. To address this limitation, we have recently developed a dynamic GSM framework using time-course metabolomics, called dynamic flux activity or DFA (Chandrasekaran et al., 2017). In our previous studies, DFA accurately predicted metabolic fluxes during transition between stem cell states, and the response of cancer cells to metabolic perturbations (Chandrasekaran et al., 2017; Nelson et al., 2019).

Here we develop a new DFA model of the IL-3-induced mammalian cell cycle using our timecourse metabolomics data. We model each cell cycle phase sequentially by DFA using our phase-specific metabolomics data, providing a comprehensive map of metabolic regulation involving all phases of the cell cycle. Our phase-specific dynamic models are validated by matching proteomics and phosphoproteomics data and also corroborate our previous findings. Furthermore, we discover phase-specific reactions and pathways that offer novel insight into biochemical mechanisms of the cell cycle.

## DATA AND METHODS

### Time-course metabolomics and proteomics data of a mammalian cell cycle

We previously generated time-course mass spectrometry-based proteomics and metabolomics data to investigate a cellular transition from a quiescence state to the first cell cycle in response to IL-3 activation in a murine pro-B cell line (Lee et al., 2017). Our proteomics, intracellular and extracellular metabolomics data sets match each other consisting of 6 time points over the first cell cycle. The proteomics was done in duplicates and the metabolomics in triplicates. To build our metabolic network model, we use the metabolomics data of 155 intracellular and 173 extracellular metabolites. We use 2,666 proteins of high confidence for model evaluation.

### Phosphoproteomics of the cell cycle

We generated phosphoproteomics data matching to our time-course metabolomics and proteomics data across (6 time points; no biological replicate; unpublished). We use the phosphoproteomics data for model evaluation as a proof-of-concept of our analytical framework in addition to the proteomics data. Briefly, phosphopeptides were enriched using titanium dioxide (TiO2) and 6-plex tandem mass tags (TMT) were used for mass spectrometry analysis as in our previous proteomics study (Lee et al., 2017). We obtained abundance profiles of high-confidence 3,095 phosphopeptides for 1,552 proteins across the 6 time points. The occurrence distribution of the 3 phosphorylated residues, serine (S), threonine (T), and tyrosine (Y), is as follows: S – 85%, T – 13%, and Y – 2%. About 96% of the 1,552 proteins have less than 6 phosphorylation sites. About 84% of them are found to be phosphorylated at one of the 3 residues, about 15% of them at 2 different residues, and 1% (14 proteins) at all of the 3 residues.

### Metabolic network data

For metabolic network modeling, we use the well-established human metabolic network reconstruction, Recon1 (Duarte et al., 2007), which is also a good approximation for mouse models as we showed in our previous work (Chandrasekaran et al., 2017). It consists of 3744 reactions, 2766 metabolites, 1496 metabolic genes, and 2004 metabolic enzymes curated from literature.

### Dynamic metabolic network modeling

We previously developed a dynamic metabolic network modeling framework, called DFA or Dynamic Flux Activity (Chandrasekaran et al., 2017), which is based on flux balance analysis (FBA) and utilizes time-course metabolomics data. In this framework, metabolite level changes over time are set to be non-zero in contrast to FBA, reflecting a non-steady state. DFA solves an optimization problem maximizing an objective, in this case, the biomass objective (*v*_biomass_) while satisfying the stoichiometric constraints that represent the metabolic network architecture. In addition, two flux activity coefficients 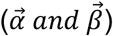 are defined for each metabolite which represent deviations from measured metabolite rate of change. DFA identifies flux vectors or distributions for all reactions in the network across the time course that best fits the metabolomics data (Fig. 1).

**Figure 1.**
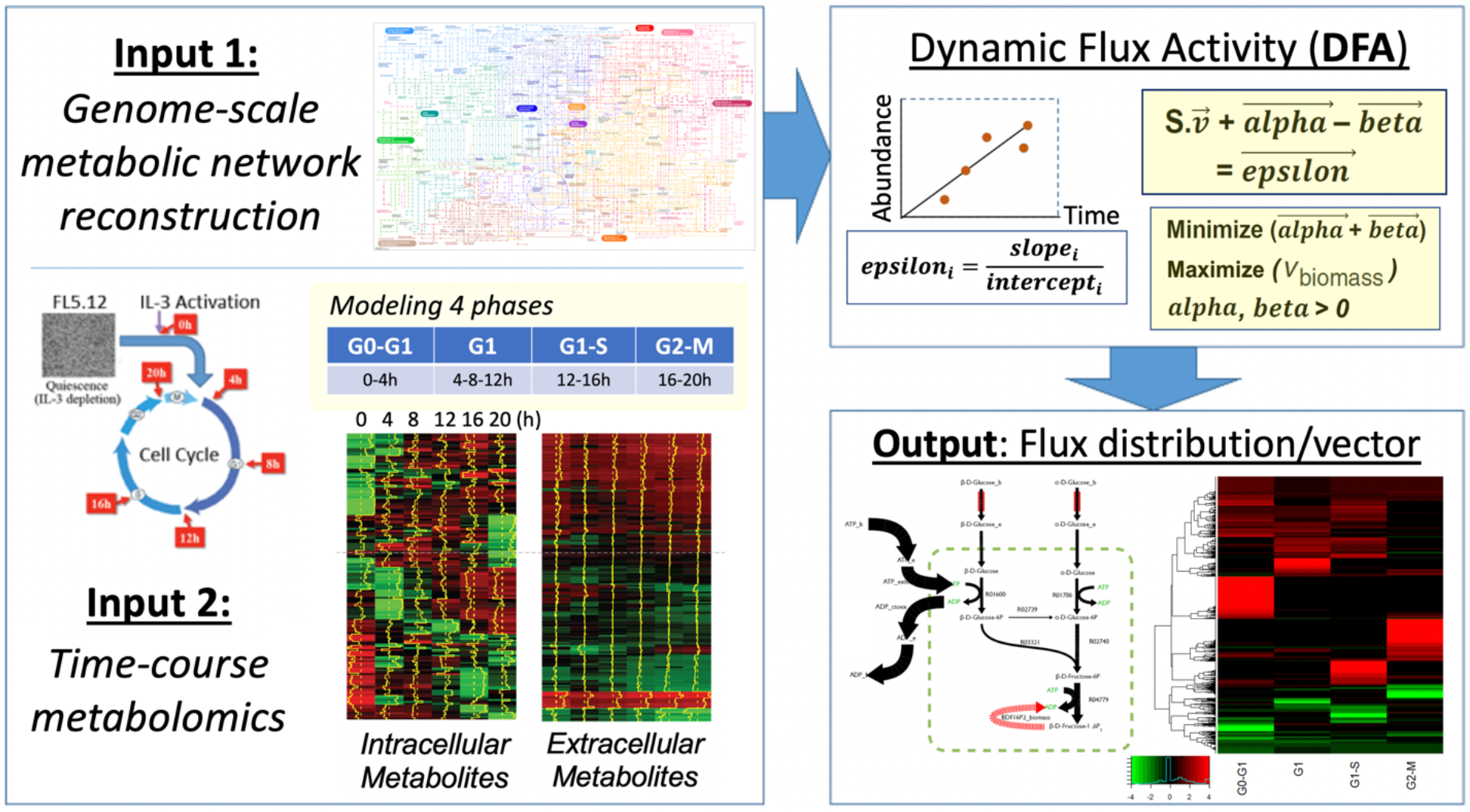
Overall workflow. We use Recon1 as our base metabolic network (Input 1; the map is from iPath3, https://pathways.embl.de/ipath3.cgi?map=metabolic) and our published timecourse metabolomics of the cell cycle (Input 2). We then build a dynamic model comprising all the 4 phases using DFA based on the time-course metabolomics data. The output is a flux distribution of all active reactions over time as shown in a toy metabolic network (taken from Wikipedia: Flux balance analysis). The heatmap shows all active reactions predicted across the cell cycle from our metabolomics data (See also Fig. 2).

To create a dynamic model of the entire cell cycle, we aim to infer metabolic flux during every temporal transition using DFA. To do this, we model the IL-3-induced cell cycle by a sequence of 4 linearized cell-cycle phases, i.e., 4 DFA models, using our intracellular and extracellular metabolomics. Since DFA requires at least 2 time points from time-course metabolomics data to calculate the relative rate of change of each metabolite, we divided our cell cycle data into 4 groups to model the cell cycle phases as follows: 0 – 4 hours for G0/G1, 4 – 12 hours for G1, 12 – 16 hours for G1-S, and 16 – 20 hours for G2/M. Therefore, we constructed 4 DFA models by mapping temporal metabolite measurements onto Recon1 and obtained flux distributions for each phase. The metabolite mapping between our metabolomics data and Recon1 was done by the MetaboAnalyst webtool (Chong et al., 2018) and manual curation using canonical IDs from HMDB and ChEBI databases and chemical formulae. The overview of our approach is shown in Fig. 1. Modeling was done in MATLAB and python as in our previous work (Shen et al., 2019b).

### Model evaluation by comparison with differentially expressed proteins and phosphoproteins

We use our matching time-course proteomics and phosphoproteomics data for evaluation of our predictions, which are orthogonal to the metabolomics data. We hypothesize that the reactions with high (or low) flux activity predicted by the cell cycle model should overlap with reactions with high (or low) enzyme levels from proteomics or phosphoproteomics data. To conduct this analysis, each proteomics enzyme was matched to its associated reaction, enabling a direct comparison between proteomics abundance and DFA model fluxes. We assess the significance of the overlap using hypergeometric tests for both top model reactions and top proteins or phosphoproteins identified using maximum fold change (MFC), standard deviation (SD), and a range of fold change thresholds.

### Model evaluation by comparison with metabolic fluxes derived from proteomics and phosphoproteomics data

We use the *linear version of the iMAT* algorithm, which we developed previously for transcriptomics-based flux prediction (Shen et al., 2019a), to predict fluxes based on proteomics and phosphoproteomics data. Fluxes from these models were then compared with DFA model predictions for further validation. To predict fluxes from proteomics data, we evaluated all 6 temporal phase pairs from the 4 phases: G0 to G1, G0 to S, G0 to G2/M, G1 to S, G1 to G2/M, and S to G2/M. For each phase pair, we used the DFA-predicted fluxes as the initial phase and the lists of up-/down-regulated proteins or phosphoproteins between the two phases to predict fluxes at the final phase. We then calculated ratios of final to initial fluxes. The same ratios of final to initial fluxes for each phase pair were computed for DFA model reactions, enabling a direct comparison between DFA reaction predictions and proteomics/phosphoproteomics reactions. To find overlaps, we used ratio thresholds of 1.1, 1.25, 1.5, and 2.0, i.e., 10%, 25%, 50%, and 100% flux increase from the initial to final phases and assessed significance with hypergeometric tests and fold enrichment.

## RESULTS

### 1. Metabolic network modeling of a mammalian cell cycle

We were able to map 126 of 155 intracellular metabolites and 131 of 173 extracellular metabolites from our metabolomics data (Lee et al., 2017) onto Recon1 (Duarte et al., 2007) for the 6 time points with 4-hour intervals covering the first cell cycle after IL-3 activation of murine pro-B cells from a G0 resting state (Fig. 1). As described in Methods, by applying DFA to the time-course metabolomics data, we constructed a sequence of 4 DFA models for 4 cell-cycle phases: 0 – 4 hours for G0/G1, 4 – 12 hours for G1, 12 – 16 hours for G1-S, and 16 – 20 hours for G2/M (Fig. 1). Using the 4 DFA models we obtained flux distributions of metabolic reactions for the IL-3-induced first cell cycle.

### 3. Flux prediction and metabolic network dynamics across the cell cycle

We predicted a total of 820 active metabolic reactions in the 4 phases of the cell cycle (Figs. 2A and 2B; Supplementary Table S1). The highest flux values in forward and reverse directions occur in the same reactions in all the phases: calcium/sodium antiporter for the forward reaction (positive flux values) and L-glutamate secretion via secretory vesicle (ATP driven) for the reverse reaction (negative flux values). The total absolute flux is highest at the G0/G1 transition and the second highest at the G1/S transition. The biomass objective is largest at the G0/G1 transition as in the total absolute flux (Fig. 2C). The second largest is in G2/M where cells divide. Flux predictions across the cell cycle phases were projected onto a map of the human metabolic network reconstruction, Recon1, for visual inspection of dynamic changes of mammalian cell cycle metabolism (Fig. 3). The highest activities across the cell cycle phases occur in different parts of the network. Nucleotide metabolism, glutathione and nicotinamide metabolism pathways have the highest activity in the G0 phase, folate metabolism in G1, glycine, serine, and threonine metabolism in G1/S, and extracellular transport in G2/M.

**Figure 2.**
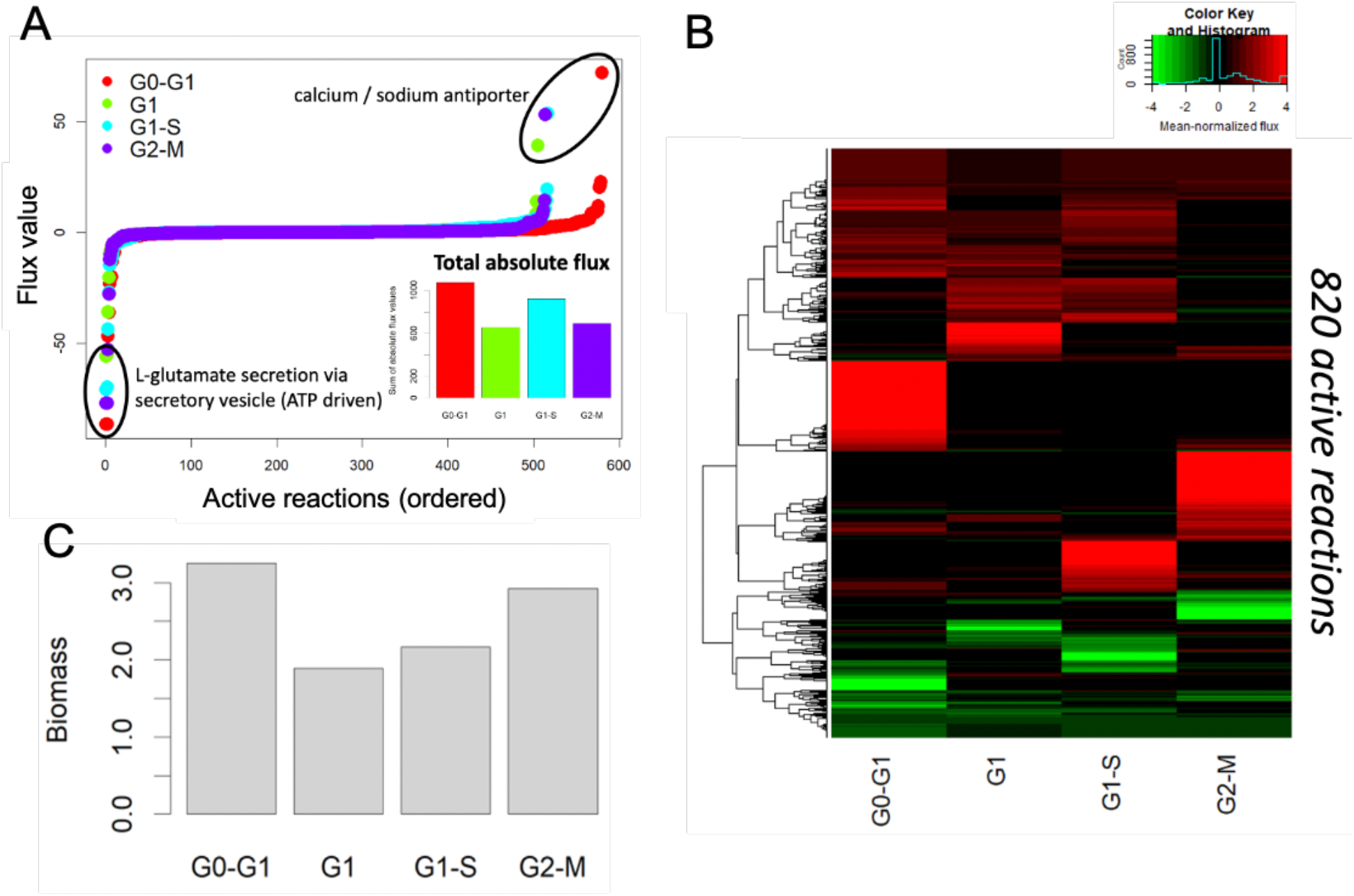
Flux prediction across the cell cycle. (A) Flux distributions for each cell cycle phase along with the total absolute flux in the inset bar plot. A total of 820 active reactions were predicted. (B) Heatmap of mean-normalized flux values across the cell cycle phases for the 820 active reactions. A hierarchical clustering was done for those reactions. (C) Optimized values of the biomass objective function across the cell cycle phases.

**Figure 3.**
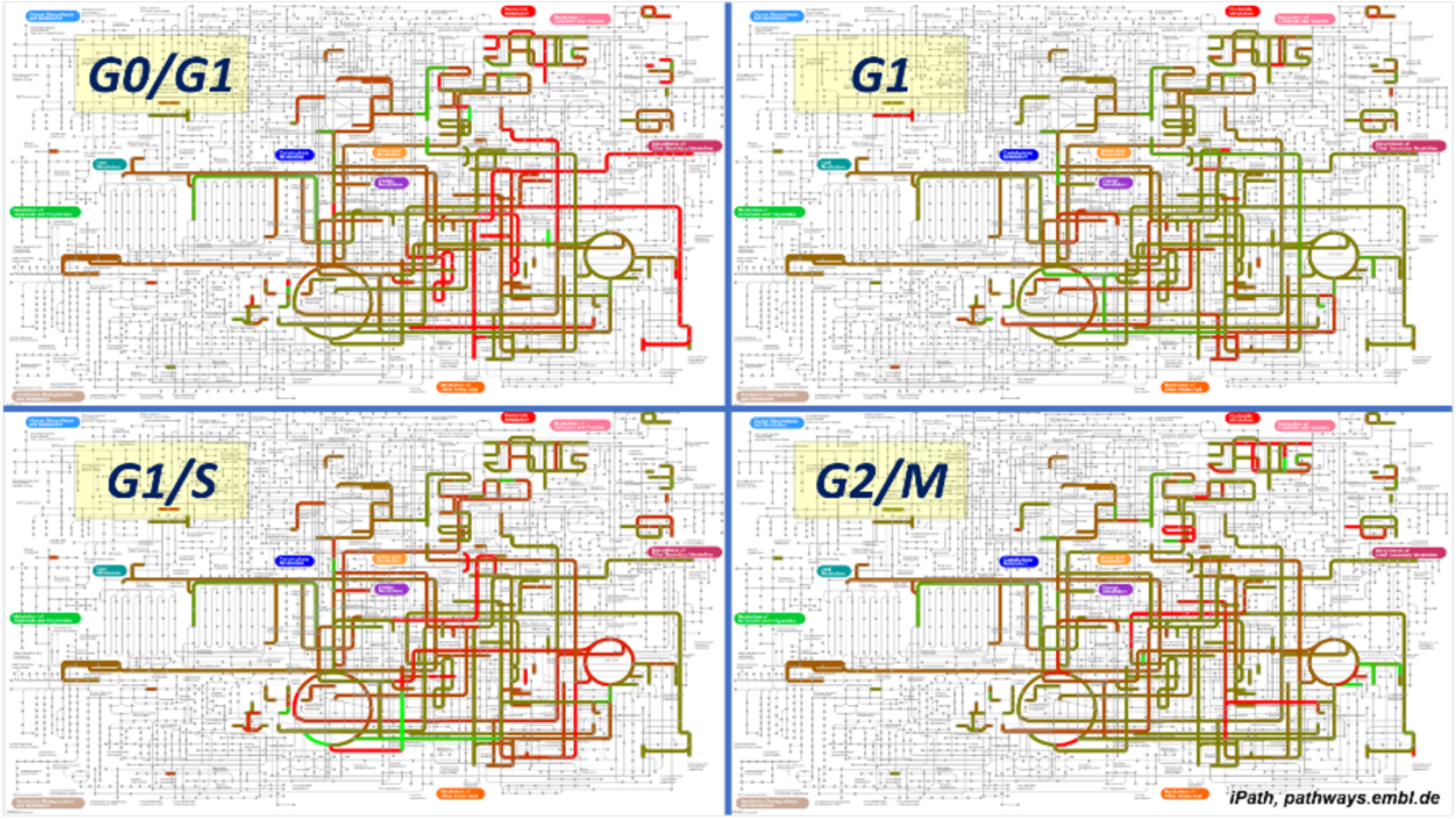
Flux prediction dynamics over the first cell cycle. Reaction flux predictions at each cell cycle phase are shown in the metabolic network map of Recon1. Red and green lines represent reactions with high flux in the forward and reverse directions, respectively, with brighter colors for higher fluxes. Grey lines are inactive reactions with no fluxes. Each network visualization was created using the iPath3 webtool (http://pathways.embl.de).

### 3. Model evaluation using proteomics and phosphoproteomics data

We first evaluated our models by comparing the top enzymes from the top reactions of the models and our matching proteomics data (Fig. 4A). All overlaps at 5 different thresholds of 10 – 50% for the top proteins from the proteomics data, either by standard deviation (SD) or by maximum fold change (MFC), are statistically significant by the hypergeometric test with p < 0.05 (Fig. 4B). This is also true when we use the phosphoproteomics data of 1,332 phosphoproteins with 3,873 phosphosites: p = 2.0e-4 ~ 0.057 for SD and p = 2.57e-6 ~ 3.3e-4 for MFC. The top phosphoproteins identify a distinct set of reactions that are not identified by the top proteins from the proteomics. For a more specific illustration of model validation, we previously discovered that methionine consumption is highest in the mid-late G1 phase of the cell cycle, among all essential amino acids (Lee et al., 2017). Our model predicted that the total flux of 9 methionine-related reactions is highest at the G1/S transition, validating our previous finding and offering insight into biochemical mechanisms.

**Figure 4.**
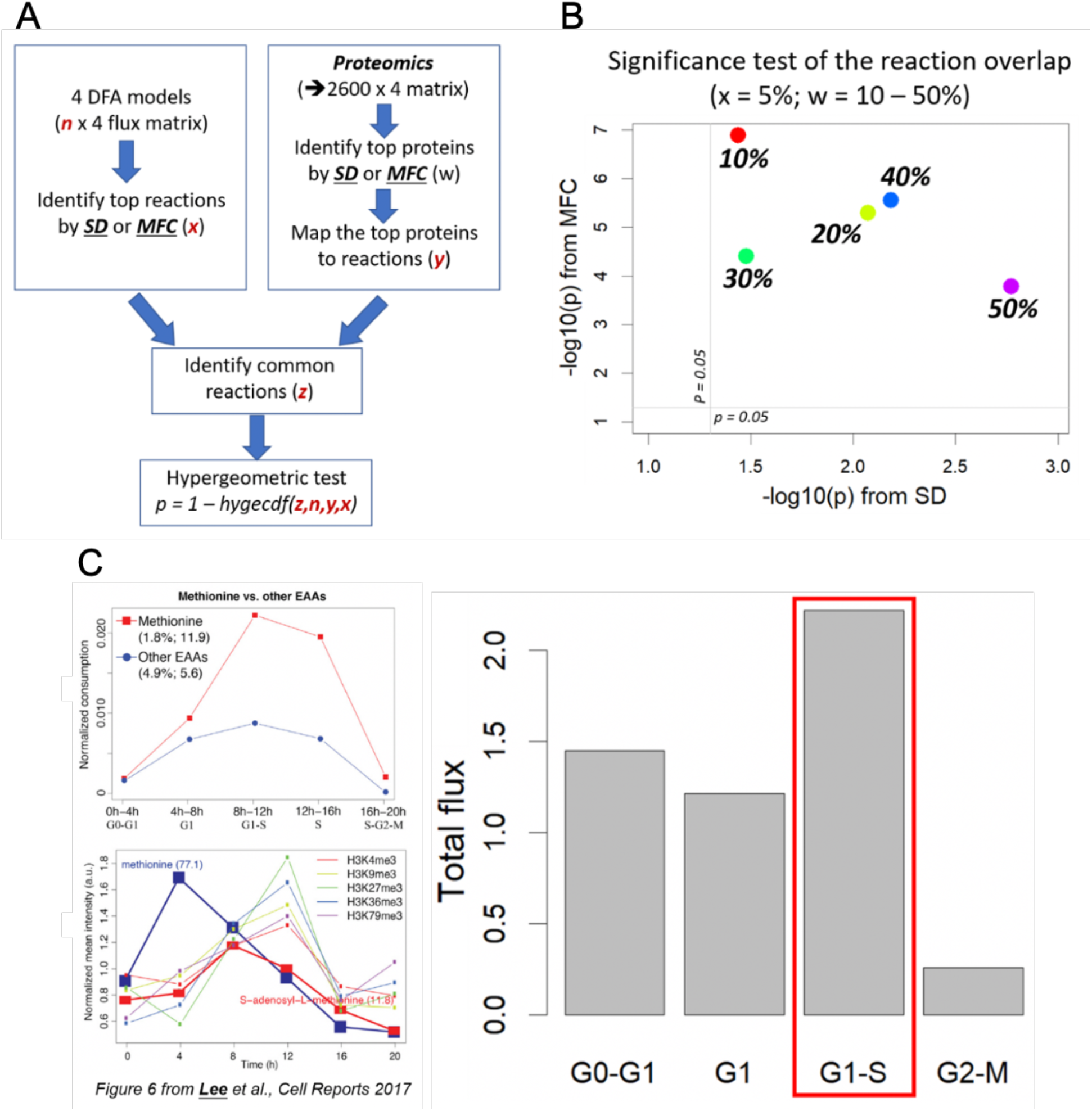
Model evaluation and validation. (A) A schematic to show how the evaluation of our models were done by comparing the top enzymes from the top reactions of the models and our matching proteomics data. (B) Hypergeometric p-values of the overlap test from (A) for multiple thresholds. (C) A model validation was done against our previous experimental data of methionine consumption and the total flux of 9 methionine-related predicted reactions.

### 4. Phase-specific reactions and processes

Having validated our model, our flux predictions reveal phase-specific reactions, which can be visualized by a heatmap of normalized flux data and a hierarchical clustering as in Fig. 5A. The number of unique forward/reverse reactions is highest at the G0/G1 transition (Fig. 5B), agreeing with the highest total absolute flux and the largest biomass objective described above (Fig. 2A and 2C). The second highest total number of unique reactions is in the G2/M phase of cell division, agreeing with the second highest biomass (Fig. 5B). Mapping from phase-specific reactions to metabolic pathways or processes also reveals phase-specific pathways and common pathways (Fig. 6). We find that lysine metabolism, nucleotides synthesis, fatty acid elongation, extracellular/mitochondrial/peroxisomal transport systems, and heme biosynthesis are major active processes at the G0/G1 transition, which suggests an inter-connected biochemical mechanism for the critical transition into cell cycle, growth, and proliferation.

**Figure 5.**
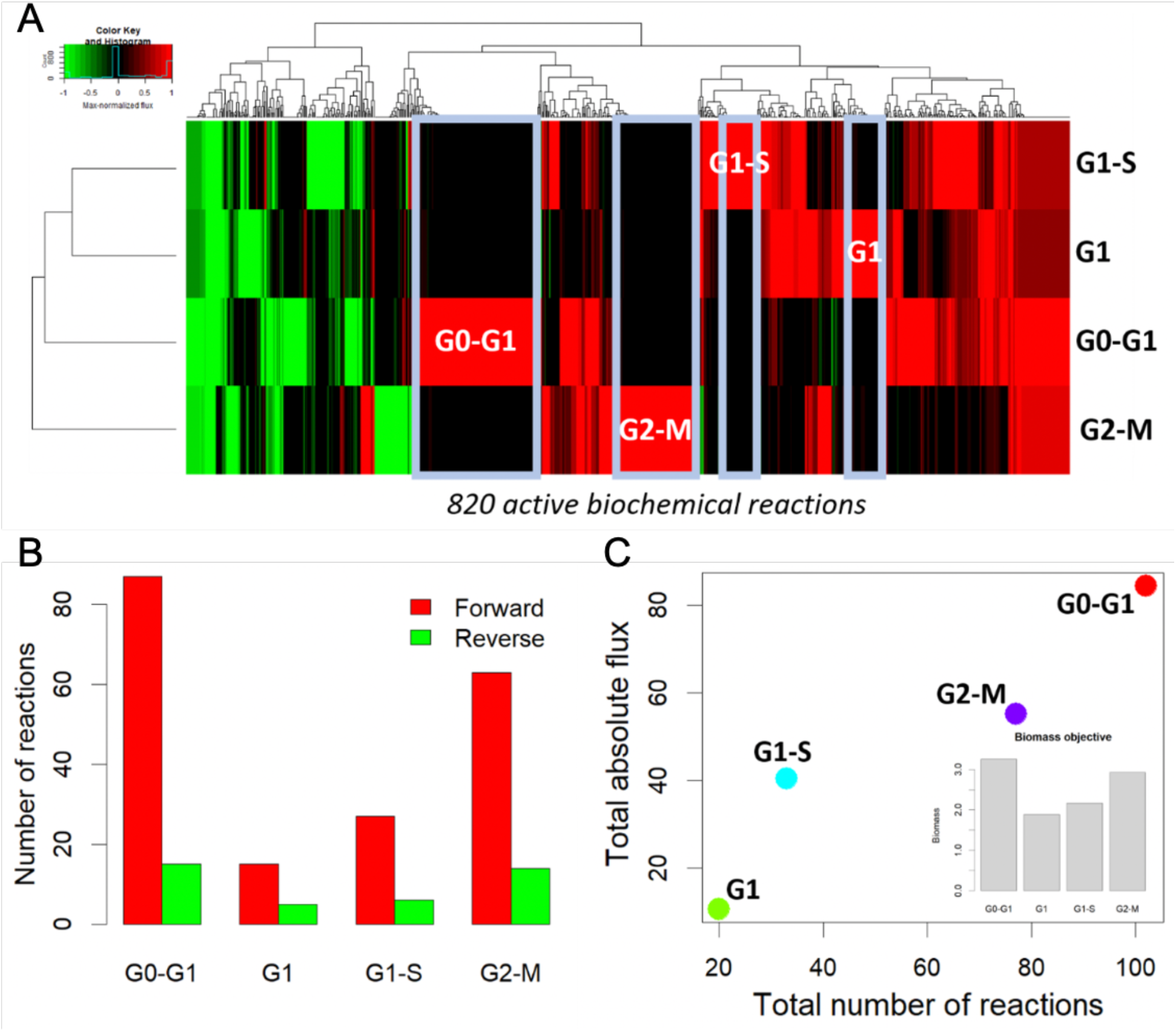
Prediction of active reactions and phase-specific reactions. (A) A heatmap of mean-normalized flux values for all 820 active reactions along with a hierarchical clustering. Phase-specific reactions are highlighted in boxes in light blue. (B) A bar plot of the numbers of phase-specific forward and reverse reactions for each phase. (C) A scatter plot of the total number of reactions and the total absolute flux for each phase. The inset is a bar plot of the biomass objective values across the phases.

**Figure 6.**
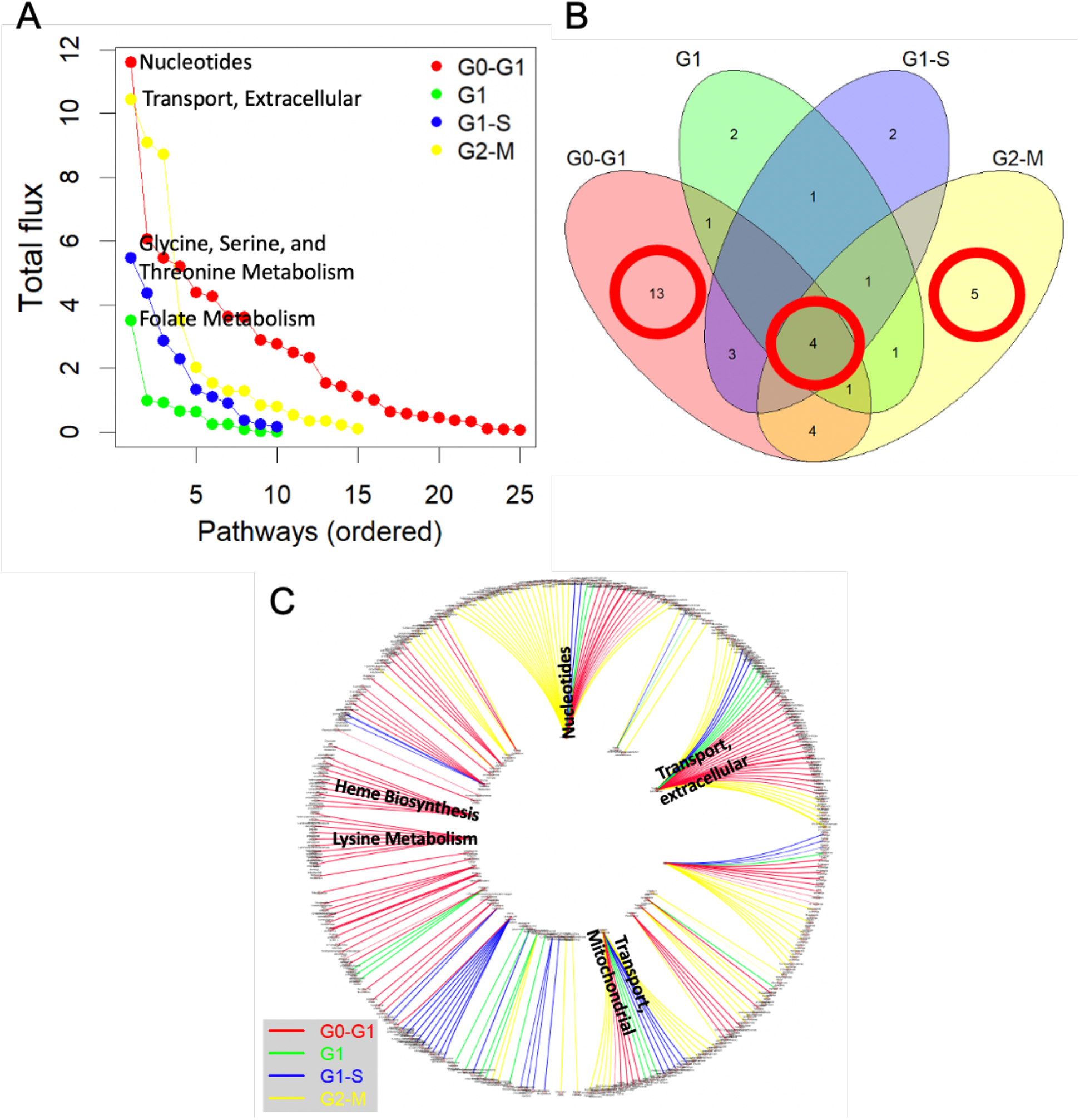
Phase-specific pathways or processes. (A) A total flux for each metabolic pathway or process based on active reactions for each phase in the decreasing order. The top pathway or process for each phase is shown. (B) A Venn diagram for pathways or processes active in individual phases. (C) A circular bipartite graph between metabolic pathways or processes (nodes on the inner circle) and reactions (nodes on the outer circle) for all phases distinguished by different colors. Some of common and phase-specific pathways or processes are shown.

### 5. Consistency with proteomics-based and phosphoproteomics-based flux predictions

With the model validation and meaningful biological interpretations above, we turned our attention to flux predictions by indirect methods based on the proteomics and phosphoproteomics data. This is to evaluate consistency with the metabolomics-based method, which is more directly related to reaction fluxes using substrate levels. To do this, DFA fluxes were used as the initial phase for each of the 6 phase pairs. Then, using a linear version of the iMAT algorithm (Shen et al., 2019a) and proteomics-derived sets of up- and down-regulated proteins between phases, fluxes for the final phase were predicted. The same approach was taken for phosphoproteomics. Fluxes from proteomics and phosphoproteomics models were compared with the metabolomics-based DFA predictions by quantifying the overlaps of top differentially active reactions for the 6 phase pairs (see Methods). At all ratio thresholds of 1.1, 1.25, 1.5, and 2.0, all overlaps are statistically significant with p-values between 6.3e-3 and 1.42e-102 by hypergeometric tests and fold enrichment values between 3.686 and 20.107 (Table 1). As an example of these overlaps, p-values at the highest threshold of 2.0 are 2.52e-89 for the G0-G1 transition and 6.26e-97 for the G0-G2. Phosphoproteomics overlaps are similarly significant with p-values of 4.52e-90 for the G0-G1 and 1.35e-93 for the G0-G2 at the 2.0 threshold. For the G0-G1 at the 2.0 threshold, 76 of the 92 proteomics fluxes overlap with model flux predictions. For phosphoproteomics at the same threshold and transition, 76 of the 91 phosphoproteomics fluxes overlap with model predictions.

**Table 1.** Consistency between metabolomics-based DFA predictions and proteomics- /phosphoproteomics-based flux predictions.

|  | Fold Change | Phase | G0 to G1 | G0 to S | G0 to G2 | G1 to S | G1 to G2 | S to G2 |
| --- | --- | --- | --- | --- | --- | --- | --- | --- |
| Proteomics | 2 | # DFA predicted | 189 | 55 | 253 | 63 | 241 | 273 |
|  |  | # proteomics predicted | 92 | 91 | 92 | 10 | 11 | 30 |
|  |  | # overlap | 76 | 13 | 85 | 3 | 5 | 14 |
|  |  | p-value | 2.52135E-89 | 2.7273E-10 | 6.25576E-97 | 0.000501027 | 0.00035545 | 4.3676E-09 |
|  |  | fold enrichment | 16.364 | 9.725 | 13.672 | 17.829 | 7.061 | 6.4 |
|  | 1.5 | # DFA predicted | 227 | 147 | 262 | 86 | 255 | 281 |
|  |  | # proteomics predicted | 100 | 95 | 100 | 15 | 15 | 93 |
|  |  | # overlap | 85 | 75 | 86 | 3 | 5 | 74 |
|  |  | p-value | 1.62858E-94 | 6.8753E-96 | 3.8955E-90 | 0.0043607 | 0.0024009 | 3.0385E-69 |
|  |  | fold enrichment | 14.019 | 20.107 | 12.289 | 8.707 | 4.894 | 10.602 |
|  | 1.25 | # DFA predicted | 233 | 165 | 268 | 104 | 265 | 282 |
|  |  | # proteomics predicted | 106 | 101 | 105 | 23 | 23 | 100 |
|  |  | # overlap | 89 | 81 | 89 | 10 | 6 | 81 |
|  |  | p-value | 4.76084E-97 | 5.9943E-100 | 3.13183E-91 | 1.49965E-91 | 0.0042941 | 3.33081E-77 |
|  |  | fold enrichment | 13.492 | 18.198 | 11.841 | 15.652 | 3.686 | 10.754 |
|  | 1.1 | # DFA predicted | 236 | 196 | 270 | 125 | 271 | 289 |
|  |  | # proteomics predicted | 110 | 105 | 109 | 24 | 24 | 102 |
|  |  | # overlap | 93 | 83 | 91 | 11 | 7 | 83 |
|  |  | p-value | 9.723E-102 | 1.8133E-93 | 8.2066E-92 | 6.39667E-11 | 0.0011415 | 1.6188E-78 |
|  |  | fold enrichment | 13.413 | 15.1 | 11.577 | 13.728 | 4.03 | 10.542 |
| Phosphoproteomics | 2 | # DFA predicted | 189 | 55 | 253 | 63 | 241 | 273 |
|  |  | # phosphoproteomics predicted | 91 | 93 | 93 | 11 | 11 | 25 |
|  |  | # overlap | 76 | 13 | 84 | 3 | 5 | 12 |
|  |  | p-value | 4.523E-90 | 3.61426E-10 | 1.3521E-93 | 0.00068066 | 0.000355451 | 3.93334E-08 |
|  |  | fold enrichment | 16.544 | 9.516 | 13.366 | 16.208 | 7.061 | 6.583 |
|  | 1.5 | # DFA predicted | 227 | 147 | 262 | 86 | 255 | 281 |
|  |  | # phosphoproteomics predicted | 95 | 101 | 101 | 17 | 16 | 93 |
|  |  | # overlap | 81 | 78 | 85 | 3 | 5 | 77 |
|  |  | p-value | 5.5517E-90 | 9.0056E-99 | 2.6317E-87 | 0.0063045 | 0.0033 | 8.5715E-75 |
|  |  | fold enrichment | 14.063 | 19.669 | 12.026 | 7.683 | 4.588 | 11.032 |
|  | 1.25 | # DFA predicted | 233 | 165 | 268 | 104 | 265 | 282 |
|  |  | # phosphoproteomics predicted | 101 | 107 | 107 | 24 | 24 | 98 |
|  |  | # overlap | 85 | 84 | 89 | 10 | 6 | 82 |
|  |  | p-value | 1.52113E-92 | 1.4208E-102 | 1.05064E-89 | 2.5119E-10 | 0.0053925 | 1.55486E-80 |
|  |  | fold enrichment | 13.523 | 17.813 | 11.62 | 15 | 3.532 | 11.109 |
|  | 1.1 | # DFA predicted | 236 | 196 | 270 | 125 | 271 | 289 |
|  |  | # phosphoproteomics predicted | 105 | 111 | 111 | 25 | 25 | 100 |
|  |  | # overlap | 89 | 86 | 91 | 11 | 7 | 84 |
|  |  | p-value | 3.20735E-97 | 9.2876E-96 | 2.3863E-90 | 1.11027E-10 | 0.001489 | 7.30189E-82 |
|  |  | fold enrichment | 13.447 | 14.8 | 11.368 | 13.179 | 3.868 | 10.882 |

Conducting the two evaluation methods above provided a more comprehensive explanation of how metabolomics-based fluxes aligned with proteomics and phosphoproteomics-derived fluxes. For example, for the G0/G1 transition at a fold change threshold of 10%, there were 236 metabolomics-based DFA reactions that were differentially active. When overlapping these reactions with the proteomics data directly, 18 of the reactions were explained. By overlapping the DFA reactions with the proteomics-based reaction fluxes, 93 reactions were explained. At the same threshold for the phosphoproteomics, 18 reactions were explained by the phosphoproteomics data directly, which were distinct from the 18 reactions from the proteomics data, and 89 were explained by phosphoproteomics-based fluxes. As such, a more comprehensive coverage of the reactions was achieved through both methods.

## DISCUSSION

Cell cycle is a fundamental process in biology. Metabolic regulation of cell cycle is an important component controlling cell phase transitions, yet its understanding has been limited by the complexity of temporal metabolic changes. Metabolic network modeling has been a major mathematical approach for studying biochemical reactions and metabolic regulation in systems biology. However, no mathematical model of mammalian cell cycle metabolism exists. Modeling metabolism of cell cycle is challenging as it involves multiple phase transitions and the metabolic effects from different omics data occur at different time scales. In addition, traditional methods for integrating omics data with metabolic network models assume a steady-state system, whereas cell cycle is dynamic. Here we developed a dynamic metabolic network model for a mammalian cell cycle using time-course metabolomics based on our previous studies (Chandrasekaran et al., 2017; Lee et al., 2017). Our metabolomics-based model can describe each cell cycle phase in terms of flux distributions, which is consistent with the matching proteomics and phosphoproteomics data. Enzymes predicted to be differentially active by the model also show changes in expression or phosphorylation levels. However, given the inherent noisiness of metabolomics data with high variability among biological replicate samples (Lee et al., 2019), our flux predictions should be considered qualitatively plausible until validated by experimental flux measurements.

Our analysis of the IL-3-induced cell cycle of murine pro-B cells revealed unique metabolic activity in each phase and that the G0/G1 transition phase has the largest number of active reactions as well as the highest biomass synthesis rate, which is meaningful because of its critical role in entry into the cell cycle and proliferation (Lee et al., 2017). In particular, there are multiple literature supports for our prediction of lysine metabolism, fatty acid elongation, and heme biosynthesis at the G0/G1 transition in relation to protein synthesis and cell growth (Gimple et al., 2019; Jang et al., 2020; Pran Babu et al., 2020; Yu et al., 2017). From the model validation using proteomics and phosphoproteomics, we find that those enzymes in our predicted high-flux reactions are likely to be enzymes of high expression changes in terms of both total proteins and phosphoproteins, as expected, and that reactions regulated by phosphoprotein level changes are distinct from those regulated by total protein level changes across the cell cycle. This may suggest a division of labor in metabolic regulation between different regulatory mechanisms, as revealed in our related work (Smith et al., 2021).

On the other hand, we showed that fluxes predicted using either proteomics or phosphoproteomics are consistent with fluxes predicted by metabolomics-based DFA predictions for all 6 phase pairs with varying thresholds. This suggests that proteomic and phosphoproteomic changes are distinct in abundance, whereas they result in flux changes in similar reactions. Of note, we point out that the overlap p-values are highly significant and several orders of magnitude higher than the overlaps observed through the first validation method. These results suggest that a small set of reactions are regulated directly by changing protein or phosphoprotein levels. By modeling systems-level metabolic consequences of regulating a small number of proteins or phosphoproteins, we were able to better explain the prediction of reaction flux changes. In addition, the predicted fluxes or reactions by the proteomics- and phosphoproteomics-based models were similar. This contrasts with the first validation case, where the reactions identified by top abundance changes from proteomics and phosphoproteomics were distinct. In this respect, the validation by the phosphoproteomics data, despite the caveat of no replicates, implies that the phosphorylation mechanism is meaningful in metabolic network modeling of the mammalian cell cycle. Indeed, independent of this work, we developed a new computational framework for predicting metabolic regulation by integrating metabolic network modeling with machine learning, called *Comparative Analysis of Regulators of Metabolism* (CAROM), to classify enzymes regulated by phosphorylation or acetylation across 4 species: bacteria, yeast, mouse, and human (Smith et al., 2021). In that study, we show that classification accuracy using our cell cycle model and the phosphoproteomics data is greater than 80%, suggesting the importance of phosphoproteomics data for mechanistic insights into metabolic regulation.

Our model is a proof-of-concept draft model and can be improved in many directions in future studies. For example, some inconsistencies in flux prediction may arise due to regulation not included in the current model such as allosteric feedback. This could be resolved by a combination of manual curation and automated curation by methods such as GEMINI (Chandrasekaran and Price, 2013). Another approach for model improvement would be Flux Variability Analysis to explore the redundancy in the network (Gudmundsson and Thiele, 2010; Mahadevan and Schilling, 2003).

In conclusion, since cell cycle is a fundamental process in biology, our model and data may have a wide range of immediate applications in understanding development, ageing, and metabolic disorders like cancer, with insights into various regulatory mechanisms that control metabolic dynamics during cell cycle and uncover phase-specific metabolic demands.

## DATA AND CODE AVAILABILITY

The processed data and flux predictions are available in Supplementary Table 1. Codes are available upon reasonable request.

## AUTHOR CONTRIBUTIONS

HL and SC conceived and designed the study. FS implemented the model. HL, FS, and AE performed the analysis. HL, FS, AE, and SC analyzed the data. HL and MPJ generated the phosphoproteomics data. HL, FS, AE, and SC wrote the paper.

## ACKNOWLEDGEMENTS

HL and MPJ would like to thank Drs. Marc W. Kirschner and Steven P. Gygi for their generous support for the phosphoproteomics experiment which was performed when they were research fellows in their laboratories. This work was supported by faculty start-up funds from the University of Michigan and R35 GM13779501 from the NIH to SC.

